# A fully open-source framework for deep learning protein real-valued distances

**DOI:** 10.1101/2020.04.26.061820

**Authors:** Badri Adhikari

## Abstract

As deep learning algorithms drive the progress in protein structure prediction, a lot remains to be studied at this emerging crossway of deep learning and protein structure prediction. Recent findings show that inter-residue distance prediction, a more granular version of the well-known contact prediction problem, is a key to predict accurate models. We believe that deep learning methods that predict these distances are still at infancy. To advance these methods and develop other novel methods, we need a small and representative dataset packaged for fast development and testing. In this work, we introduce Protein Distance Net (PDNET), a dataset derived from the widely used DeepCov dataset and consists of 3456 representative protein chains for training and validation. It is packaged with all the scripts that were used to curate the dataset, generate the input features and distance maps, and scripts with deep learning models to train, validate and test. Deep learning models can also be trained and tested in a web browser using free platforms such as Google Colab. We discuss how this dataset can be used to predict contacts, distance intervals, and real-valued distances (in Å) by designing regression models. All scripts, training data, deep learning code for training, validation, and testing, and Python notebooks are available at https://github.com/ba-lab/pdnet/.

## 1 Introduction

Deep learning and covariance signals obtained from sequence alignments are speeding up the progress in the field of protein structure prediction [1]. It is exciting that information culled from sequences whose structures are not solved can serve as the primary input feature to predict contacts and distances. The most successful methods in the recent CASP13 experiment (http://predictioncenter.org/), a biannual protein structure and contact prediction competition, exploit such sequence databases and have unanimously demonstrated that a key to push the current progress is accurate contact and distance map prediction [2, 3, 4]. The distance prediction methods, in particular, are a major advancement in the area of ab initio or free modelling. We believe that a lot remains to be investigated at this emerging cross-way of deep learning and protein structure prediction. For example, we do not know if current deep learning methods are an ideal fit for solving the distance prediction problem. Do we need novel algorithms specifically designed for structure prediction? In addition, many kinds and combinations of input features such as secondary structure predictions, coevolution-based signals [5], raw features such as pair frequencies matrix, covariance matrix [6], compressed covariance matrix [7], and precision matrix [8] are currently used as input features. How to best engineer these features for deep learning algorithms also remains an open question. An irony about current methods for structure prediction is that they rely on coevolution and conservation signals in multiple sequence alignments which an amino acid sequence, folding in a cell, does not have access to. Hence, there is some possibility that we can build accurate models without such input features. How accurately we can do this also remains an open question. We believe that there is an urgent need for a small and representative dataset packaged for fast development and investigation, and PDNET attempts to fill this gap.

The protein inter-residue distance prediction problem is to predict a pair-wise distance matrix (2D) from a protein sequence (one-dimensional sequence of amino acids). It may be compared with the monocular or stereo depth estimation problem in computer vision (see Figure 1). Similar to a depth prediction problem [9], the input is a three-dimensional volume (height × width × channels) and the output is the same dimensions as the input (height × width). Many characteristics of distance prediction, however, make it a unique deep learning problem. Unlike problems in computer vision which usually have one to three input channels, the distance prediction problem may have a few to a few hundred input channels depending on what and how many input features are used. Also a distance map is symmetrical along the diagonal and each pixel on the map represents a distance between a pair of residues in the sequence. A single input feature, such as a precision matrix, can have more than 400 channels. Some of these input channels are tiled input rows or columns obtained from one-dimensional predictions such as secondary structures and solvent accessibility. The comparison of this problem with a computer vision problem naturally raises the question of what the convolutional filters in each layer learn. Visualization of what the deep learning layers learn and how the input features translate over the layers is difficult and only some research has been done in this area [10]. Because we do not know what to expect in our visualizations, exploiting the techniques of explainable artificial intelligence used in computer vision are not directly useful for the distance prediction problem. Another unique feature of the distance prediction problem, compared to computer vision problems, is that a protein structure and the corresponding distance maps cannot be augmented in ways that images can be. For instance distance maps cannot be scaled, rotated, or flipped. This is because an object in the real world (for example, a chair) may be tiny or large but in the case of protein structures, all proteins, small or large, are comprised of a fixed size structural units such as an alpha helix. In a distance map, distance pixels away from the diagonal are more informative for reconstructing the original structure. These distances, known as long-range distances, are also harder to predict (see Figure 2). In a way, this is similar to saying that in the case of depth prediction, it is more important to predict the depth of objects closer (to the camera) in the scene than the objects far away. An ideal distance prediction algorithm should predict exact physical distances in the entire distance map accurately. This is too difficult. Hence, a binary version of the problem known as contact-prediction was more widely studied.

**Figure 1:**
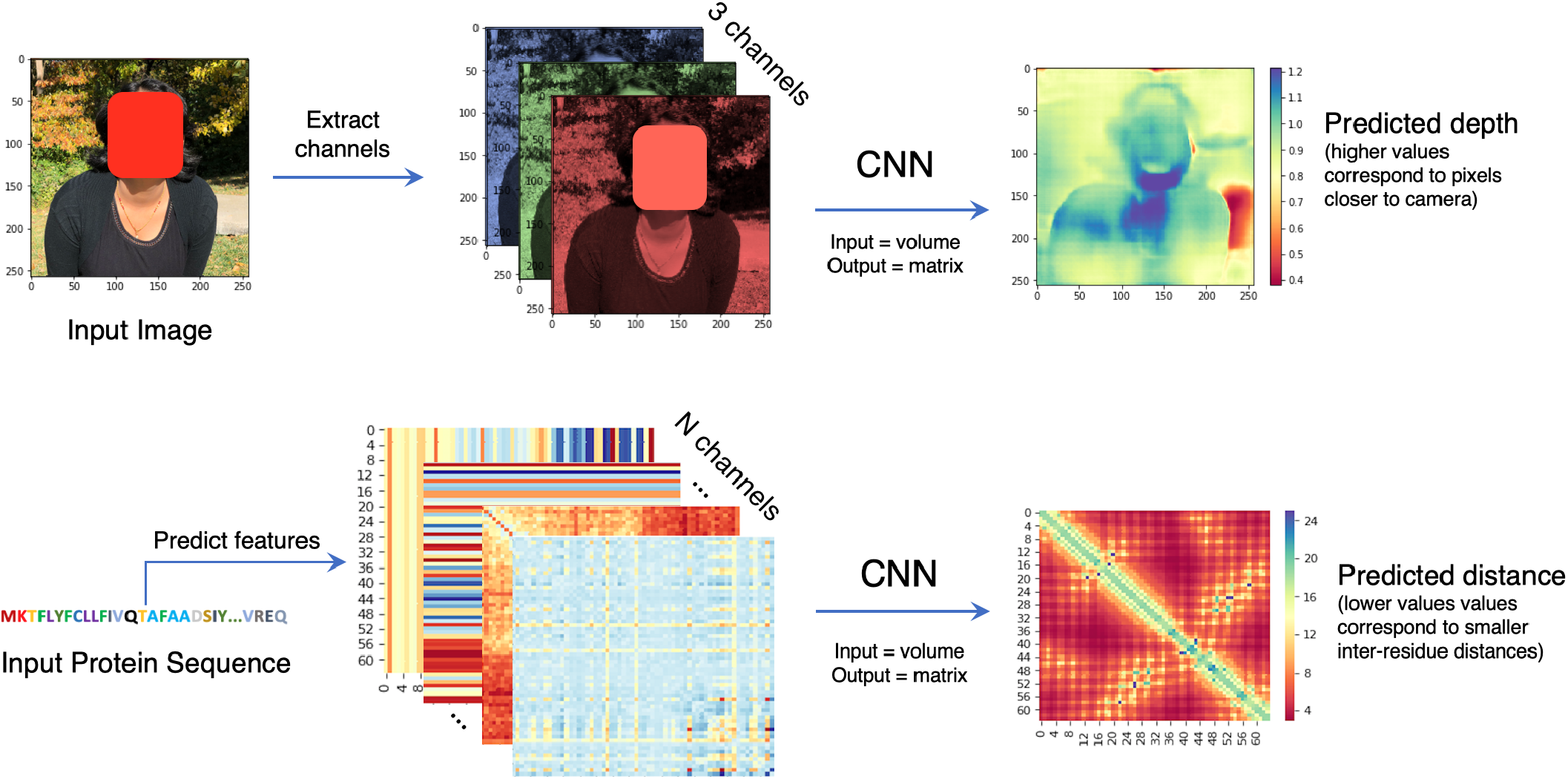
Comparison of the protein inter-residue distance prediction problem with the ‘depth prediction from single image problem’ in computer vision. In both problems the input to the deep learning model is a volume and the output is a 2D matrix. The depth predictions for this specific image (top right corner) were obtained by running the pretrained FCRN method [9]. Face in the figure is obscured because of bioRxiv policy.

**Figure 2:**
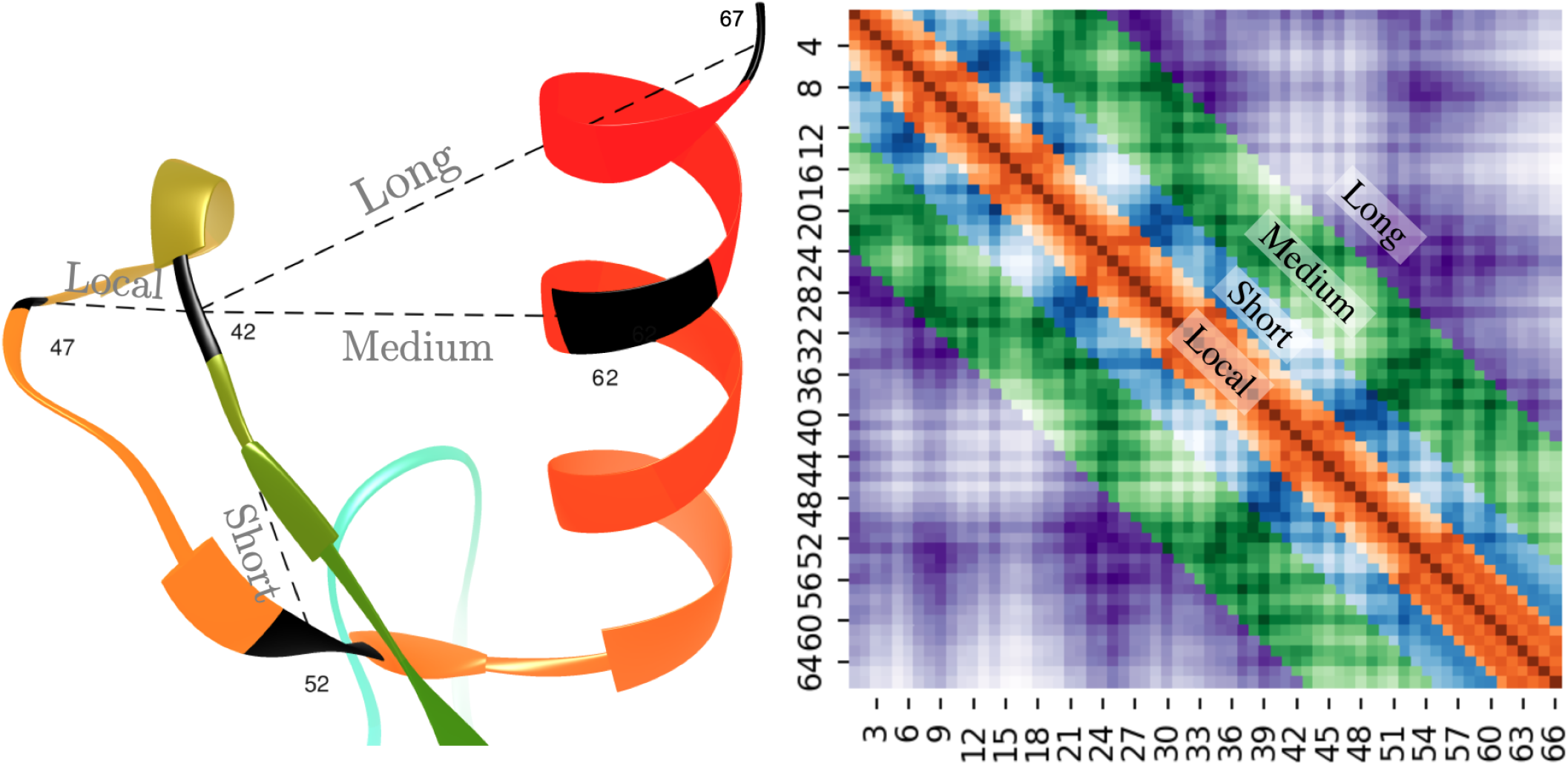
Example of long-range, medium-range, short-range, and local distances in a protein distance map. The distances between residue pairs 42-47, 42-52, 42-62, and 42-67 are examples of local, short, medium, and long-range contacts, respectively (left). In the heatmap plot (right) the sequence separation domain for long, medium, and short-range distances and local distances are [24+], [12, 23], [6, 11], and [0, 5], respectively. The labels in the x and y axis refer to the residue index in the corresponding protein sequence. The diagonal line shows that the residue pair i and i have a zero distance.

Although almost all currently successful methods [2, 3, 11] formulate the problem as a multi-class classification problem, undeniably, the ultimate goal remains to accurately predict real valued distances. Recently, some have formulated the problem as a regression problem and demonstrated promising results [12, 13].

## 2 MATERIALS AND METHODS

### 2.1 Dataset Preparation

Our primary data set is derived from the 3456 representative protein sequences and 150 test protein chains used to train, validate, and test the widely used DeepCov method [14] for protein contact prediction. The 3456 protein chains used for training and validating the DeepCov method are a subset of a set of 6729 protein chains that have no domains homologous with the protein chains in the test set. The sequence lengths of the protein chains range from 50 to 500 residues and 50 to 266 residues in the training set and test set, respectively. The original set of 6729 protein chains have less than 25% pairwise sequence similarity and hence are representative of the PDB set. We use this list of 3456 protein chains, the list of 150 protein chains, and the corresponding alignments as the input to prepare our dataset. For each fasta file in the DeepCov set, we downloaded the corresponding protein chain from www.pdb.org and cleaned it by removing rows containing non-standard amino acids and renumbered the residues to match the fasta sequence. Pairwise *Cβ* distances between residues are calculated from the true structure (3D) to obtain a distance map (2D). The amino acid sequence obtained from the true structure is used as input to predict various sequence features (1D) and pairwise features (2D), which are all translated to an input volume (2D with channels). This input volume is the input to a deep learning model and the output labels are the true distances. For maintaining consistency with DeepCov method, in all our experiments, we select the first 130 chains when sorted alphabetically by PDB IDs (a random subset) in the entire DeepCov set as the validation set and the remaining 3326 chains as the training set. The alphabetical ordering ensures a random selection because PDB IDs are automatically assigned and do not have meaning [15]. Since the 150 proteins in the test set are easier (i.e. they have much richer alignments), we further test our methods on two hard datasets - 131 CAMEO-HARD (https://www.cameo3d.org/) targets released between December 8, 2018 and June 1, 2019 and the CASP13 free-modeling targets released in 2018. These datasets are also used by other successful methods such as trRosetta [16] to evaluate the performance of their methods. Both of these datasets serve as test set as they were released much later after the original DeepCov set (training set) was curated.

It is worth noting that our dataset may not be a fully balanced representation of all the protein structural folds as summarized in the databases such as CATH [17] but it is a represenative set of the protein sequence space. Designing and developing state-of-the-art models may require a much larger dataset. However, to answer many fundamental questions about the applicability and limitations of deep learning to solve the distance prediction problem, we believe that a small dataset such as PDNET will work well.

### 2.2 Input Features

Successful methods for contact and distance prediction use a variety of features predicted and derived from the input sequence. These include predicted secondary structures, coevolution features, solvent accessibility, position-specific scoring matrix derived features, Atchley factors, many pre-computed statistical potentials, alignment statistics such as the number of effective sequences, Shannon entropy sum, mean contact potential, normalized mutual information, covariance matrices, precision matrices, etc. Contact and distance prediction methods use a variety of combinations of these features and we do not fully understand, in general, which of them contain complementary information and which are redundant. Additionally, generating all of these features for an input protein sequence is computationally expensive and takes time. Based on the findings of other recent works [18, 19, 16], we identified the top seven features that are complementary and most informative - sequence profiles, secondary structure predictions and solvent accessibility predictions using PSIPRED [20], coevolutionary signals predicted using CCMpred [5] and FreeContact [21], contact potentials calculated from multiple sequence alignments, and Shannon entropy of the alignment column. The secondary structure predictions by PSIPRED are probabilities of each residue in the input sequence being a helix, strand, or coil, i.e. predicting whether each amino acid will be a part of a helix, beta-strand or a coil in the final model. Similarly, the solvent accessibility predictions by PSIPRED are binary predictions of hydrophobicity for each residue, i.e. predicting whether each amino acid will be ‘exposed’ to water or not. Looking into the predicted multiple sequence alignment, the coevolutionary predictions by CCMpred and FreeContact capture the strength of covariance between all pairs of residue positions. These predictions themselves can be considered as contact predictions. However, deep learning algorithms can improve these predictions by learning to detect noise, correct mistakes, and identify high-confidence and missing predictions [22]. With the predicted multiple sequence alignment as input, we also calculate a contact potential matrix and the Shannon entropy sum at each residue position using the ‘alnstat’ program in the MetaPSICOV package [23]. The potential matrix captures the frequencies of the co-varying pair positions weighted by the value of each sequence in the input sequence alignment, and the Shannon entropy sum calculates the variability at each residue position. Finally, we also generated sequence profiles from the multiple sequence alignments as our last input feature.

Features derived directly from the MSA such as covariance or precision matrix are significantly important, with at least a few percent expected improvements [7]. However, in this work, we intentionally skip investigating such features because working with these features requires huge storage capacities, solid state disks, and many high-end GPUs. In a separate work, we will elaborately discuss the importance of all of these features (a work in progress).

### 2.3 Evaluation of predicted distance maps

A primary goal of evaluating the distances in a predicted distance map is to assess their usefulness towards building full three-dimensional models. We use two sets of metrics for evaluating predicted distances - a) mean absolute error (MAE) of predicted distances, and b) precision of the contacts derived from the predicted distances. To calculate MAE, we first keep all the true distances below a certain distance threshold from the native structure, and calculate mean absolute difference between these true distances and corresponding predicted distances. Analogous to the definition of various types of contacts, we define local, short-range, medium-range, and long-range distances as the distances between residue pairs with sequence separation of [0, 5], [6, 11], [11, 23], and [23+] respectively. Previous studies have shown that long-range contacts, i.e. pairs separated by at least 23 residues in the sequence, are the most informative pairs for accurate reconstruction [24, 25]. Hence, we designed our evaluation metrics focusing on long-range distances (see Figure 2). Here we evaluate the mean absolute error of medium and long-range and long-range only distances at distance thresholds of 8Å and 12Å (MAE_8_ and MAE_12_). We believe that other variations of MAE, such as root mean squared deviation (RMSD) may also be relevant. When evaluating the predictions for the validation set, we observed a Pearson’s correlation coefficient of 0.9 between MAE_8_ and P_NC_.

Ideally, to obtain contacts from predicted distances one would simply consider distances less than 8Å as contacts. Such techniques, however, did not favor the evaluation of predicted distances. Hence, we resort to a technique similar to the one that is currently used for evaluating predicted contacts. We assign contact scores such that shorter predicted distances have higher scores than longer ones and a score of 0.5 is assigned for a predicted distance of 8Å. Precisely, if *D_ij_* is the predicted distance between two residues i and j, then the corresponding score *P_ij_* is given by,

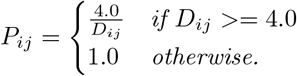

Following the practice to evaluate predicted contacts by calculating the precision of top L/x contacts, we evaluate top L, and top NC long-range contacts (*P_L_* and *P_NC_*). Here L and NC stand for the length of the protein sequence and the total number of contacts in the native structure. We calculate precision as the ratio of the number of matches and the total number of contacts considered. An exception is that - for an input protein sequence, if the corresponding true contact map does not have any long-range contacts, we exclude the target from evaluation. Our motivation for evaluating the precision of top NC contacts is driven by two insights. In a recent work [26] the Cheng group have reported that selecting or evaluating top L/x contacts does not work well for all proteins. Also, in the most recent CASP competition, the accessors of the contact prediction category have discussed many reasons for considering to evaluate top NC contacts instead of fewer contacts [25]. Although the *P_NC_* metric is not discussed for most contact prediction method papers, we believe that as more and more accurate contact prediction methods are being developed, this will emerge as a more informative and reliable metric. Table 1 summarizes our evaluation metrics.

**Table 1:** Evaluation metrics used for evaluation of predicted distances and/or contacts. L stands for the length of the protein sequence in the native (true) structure and NC is the number of true contacts in the structure.

| Prediction | Metric | Description |
| --- | --- | --- |
| Distances | $MAE_8$ | MAE of the predicted long-range distances whose corresponding true distances are shorter than 8 Å |
| | $MAE_{12}$ | MAE of the predicted long-range distances whose corresponding true distances are shorter than 12 Å |
| Contacts | $P_L$ | Precision of top L long-range predicted contacts |
| | $P_{NC}$ | Precision of top NC long-range predicted contacts |

### 2.4 Residual network architecture

All successful methods for contact and distance prediction use residual networks and their variants [3, 11, 2, 4, 16]. We developed deep-learning methods to predict contacts (DL-Contact), distance intervals (DL-Binned), and real-valued distances (DL-Distance), i.e. three separate models. Our deep learning architecture for contact prediction is a standard 128 block residual neural network with Dropouts added in between the convolutional layers as suggested in the DEEPCON method [19]. Each residual block consists of the following layers - batch normalization, ReLU activation, 2D convolution using 3×3 filters, a dropout layer with the dropout rate of 0.3, another ReLU activation, and another similar 2D convolution layer. Each network has around 9.5 million parameters. For our DL-Contact method, we set the last layer’s activation as ‘sigmoid’ and loss is calculated using binary cross-entropy. Similarly, for DL-Binned, the last layer’s activation is ‘softmax’ and loss is calculated using categorical cross-entropy, and for DL-Distance the last layer’s activation is left as ReLU. We train a model for up to 64 epochs, and during each epoch of training, we randomly crop the input feature volumes to a 128 × 128 sub-volume. Before cropping, we also pad zeros of width 5 to all sides of the input volume similar to AlphaFold’s approach [3]. During prediction, however, we build a model as big as the input sequence, i.e. we do not crop during prediction, load pretrained weights, and then predict.

### 2.5 Binned distance prediction

Considering that shorter distances are more critical for structure reconstruction and other uses, we binned our distances so that the bins are narrower for shorter distances and wider for larger distances. Specifically, we used a fixed bin width of 0.2Å for bins below 8Å and increasing bin width for larger distances (by a factor of adding 0.2Å for every next bin). These threshold for distance bins are 4, 4.2, 4.4,…, 8, 8.4, 9.0, 9.8, 10.8, 12,…, 21, 23.4, 26, and 26+Å. Our technique is slightly different from a fixed width binning technique used in methods such as RaptorX [11] and AlphaFold [3], and the two-width binning used in the DMPfold method [27]. We use the standard categorical cross-entropy loss to optimize our model with softmax output as the last layer’s activation.

To translate the predictions into contact prediction probabilities, we sum all the probabilities in the bins below the 8 Å threshold. Similarly, to translate the predictions into distance maps, for each residue pair, we pick the distance bin with the highest probability and calculate the mean of the distance range as the predicted distance. For instance, if the highest probability bin for a residue pair is [6.5, 7], then the predicted distance becomes 6.75 Å.

### 2.6 Real-valued distance map prediction

Predicting continuous distance values, like many other regression problems, is challenging. Considering the distance prediction as a regression problem, in particular, has a unique domain specific characteristic - it is more important to predict shorter distances more accurately than longer distances. This is because, from the perspective of structure prediction and binding-site prediction, it is more meaningful to predict inter-residue interactions than non-interactions (i.e. closer/smaller distances are more important). The Fisher’s group refer to such interactions as ‘interaction hubs’ [25]. It may immediately follow that the commonly used machine learning loss functions such as mean squared error or mean absolute error are unfit for this purpose because they focus on optimizing the longer distance values before the shorter ones. Recently, the ‘logcosh’ loss (logarithm of hyperbolic of cosine) is found to be highly effective for many problems. It behaves similar to the squared loss for smaller loss values and similar to absolute loss otherwise, i.e. the loss is not so strongly affected by the occasional incorrect predictions. However, this loss function also does not focus on optimizing the shorter distances. As a solution, we propose a novel loss function that precisely focuses on optimizing the shorter distances first. The idea is to invert (reciprocate) the true and predicted distances separately and then apply the logcosh loss to the difference of the two. This ‘inverse’ log cosh loss is given by

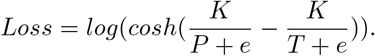

Here, P is predicted distance matrix, T is true distance matrix, e is a small positive number (epsilon), and K is a scalar that scales the losses to prevent underflow. We empirically set K to 100. Our initial implementation of this loss function, in Tensorflow, however, did not converge fast. We do not fully understand why such a loss function is not very efficient when implemented. As a solution, we inverted our distance matrices (output labels for the deep learning model) instead of inverting the loss function, i.e. we invert the input distance matrices and use the standard logcosh loss (see Figure 3). Such a setting, we find, makes it easier for the deep learning setup to converge reliably.

**Figure 3:**
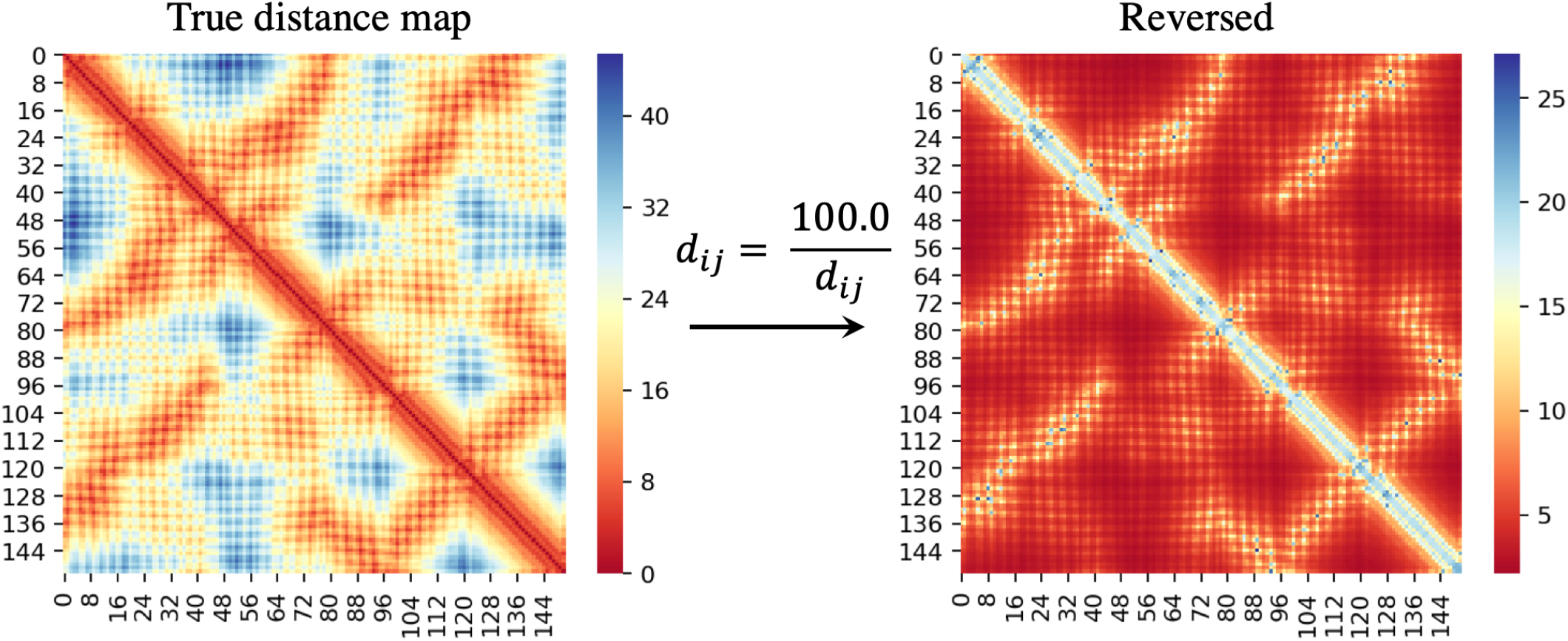
Inverting (reciprocating) the distance matrix so that larger numbers represent shorter distances in the input distance matrix. The input distance matrix is shown in left and the inverted distance matrix on the right. To avoid division-by-zero errors, all the diagonals in the input distance matrix are replaced by the mean of their neighbors.

## 3 RESULTS AND DISCUSSION

### 3.1 Dataset

PDNET, when zipped, is only around 1GB in size. All the scripts used to curate the dataset, generate the input features and distance maps, and scripts with deep learning architectures for training, validation and testing are released along with the data.

### 3.2 Evaluation of predicted distances and contacts

Here we present the evaluation of our deep-learning methods that predict contacts (DL-Contact), distance intervals (DL-Binned), and real-valued distances (DL-Distance), on the 150 proteins in the test set. Overall, we find that all methods perform relatively well on the test set because of the large and high-quality input alignments in the set (see Table 2). *P_L_* and *P_NC_* for the DL-Contact method are 69.5% and 61.1%, respectively. It is worth noting that these precision values cannot be directly compared with the precision of DeepCov [14], DEEPCON [19], or the Yang group’s methods [7] because they use covariance matrix as input features (not sequence features). However, the precision values we achieve are comparable to the results of these methods because we use the same alignments to generate our input features.

**Table 2:** MAE and precision of contact prediction method (DL-Contact), distance prediction method (DL-Distance), and binned distance prediction method (DL-Binned), on the 150 proteins in the test set. All MAE are in Å and precision are in percentage.

| Method | $MAE_8$ | $P_L$ | $P_{NC}$ |
| --- | --- | --- | --- |
| DL-Contact | - | 69.5 | 61.1 |
| DL-Distance | 4.1 | 67.5 | 59.1 |
| DL-Binned | 3.8 | 67.8 | 60.5 |

The contact precision of our multi-class classification method, DL-Binned, is lower than our binary predictor. *P_NC_* for DL-Binned and DL-Contact are 60.5% and 61.1%, respectively, on the test dataset (see Table 2). These findings slightly contradict with the findings of the Xu group [28] where they have found up to 4% improvement with their method based on binning. We believe that this slightly poor performance of a multi-class classification model on the mean absolute error and precision metrics is compensated by the predicted probability information for each class which can be used to build distogram potentials [3] for building models. We also believe that our architecture, hyper-parameters, and overall training techniques can be improved to achieve much better performance on these metrics. For instance, we found that binning in a way such that more classes are below the 8Å threshold improves contact prediction precision.

Our DL-Distance method predicts long-range distances with an MAE_8_ of 4.1Å on the test set. It is worth noting that this value seems higher because we MAE_8_ is the evaluation of all true distances below 8Å, i.e. some incorrect predictions can significantly impact the average error. Results in Table 2 show that DL-Distance, in general, predicts contacts with slightly less precision than DL-Contact. *P_NC_* of DL-Distance is around 3.3% less than that of DL-Contact. The DL-Distance method, however, predicts granular information contained in real-valued distance predictions which can be potentially more informative than the binary contact/non-contact prediction. As an example, in Figure 4 we visualize and compare the predicted contacts, binned distances, and real-value distances predicted by our three methods, for the protein chain 1a6mA in the test dataset.

**Figure 4:**
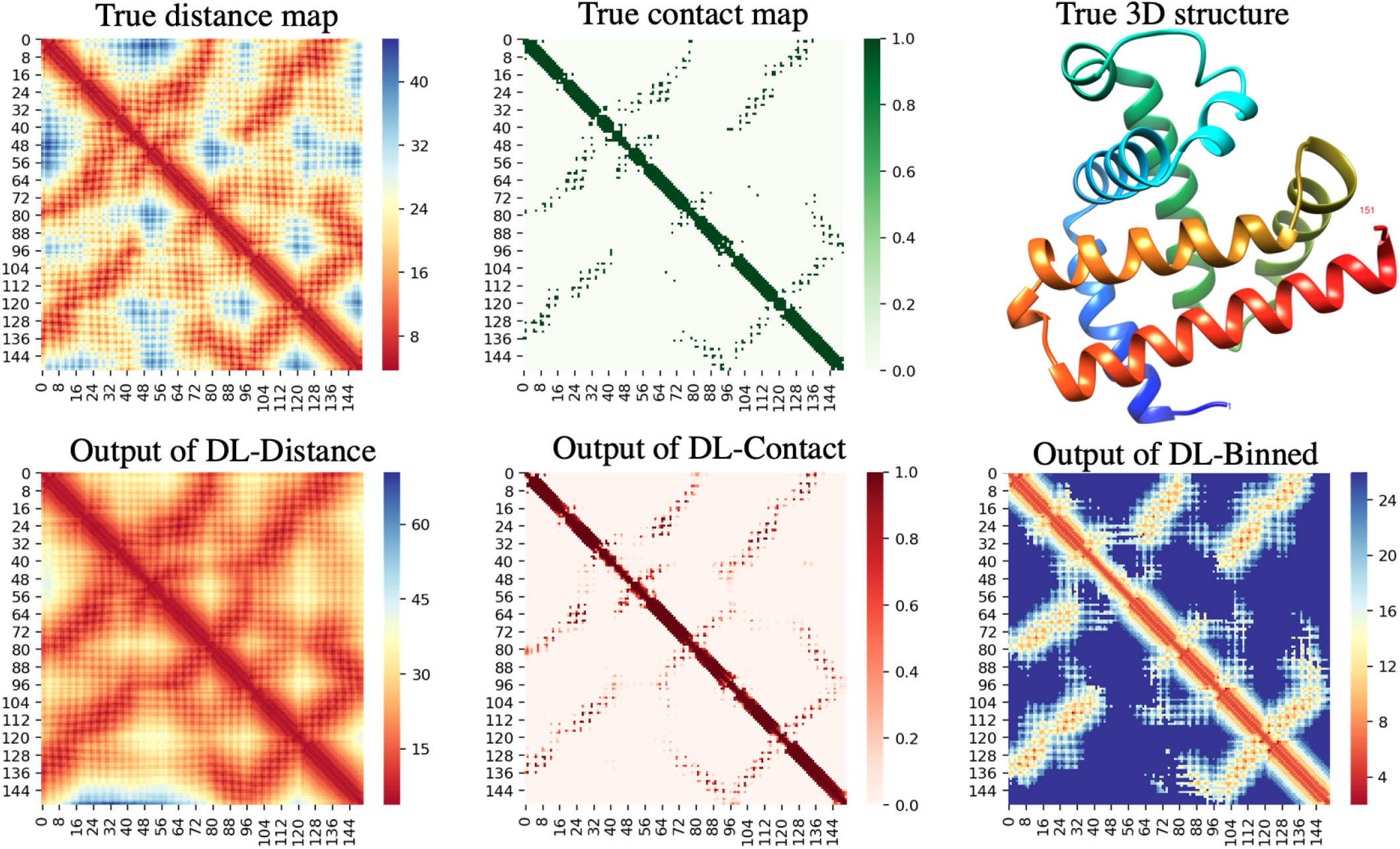
True distance map, true contact map and the native structure of chain A of the protein ‘1a6m’ in the test set (top row). Output of DL-Distance, DL-Contact, and DL-Binned (bottom row).

We also trained our model to predict inter-residue *Cα* (carbon-alpha) distances in addition to *Cβ*, and we did not find significant improvement in mean absolute error or contact precision. This can explain why methods such as the ones developed by the Xu group use separate models to predict *Cα – Cα* or *Cγ – Cγ* distances [28]. Furthermore, our DL-Binned method was extremely slow to train. On average, for one epoch of training with a batch size of 2, our DL-Binned method took around 10 hours to train, compared to our DL-Contact and DL-Distance which only took 20 minutes. In general, we observed that the training time of our multi-class classifier was proportional to the number of classes or distance bins.

### 3.3 Real-valued distance predictions

Our real-valued distance prediction method (DL-Distance), trained by inverting (reciprocating) the distance maps, helps the deep learning optimization focus on correctly predicting shorter distances before optimizing the longer ones. In other words, the model first attempts to predict shorter distances (residue pairs that are closer in physical distance, not sequence separation) over others. To visually investigate the model’s focus on shorter distances, we randomly selected two proteins from the test set, and plotted the predicted long-range distances vs the true long-range distances. Figure 5 shows that the model makes more precise predictions for shorter distances. These visualized examples validate that our method effectively focuses on correctly predicting the shortest distances over others.

**Figure 5:**
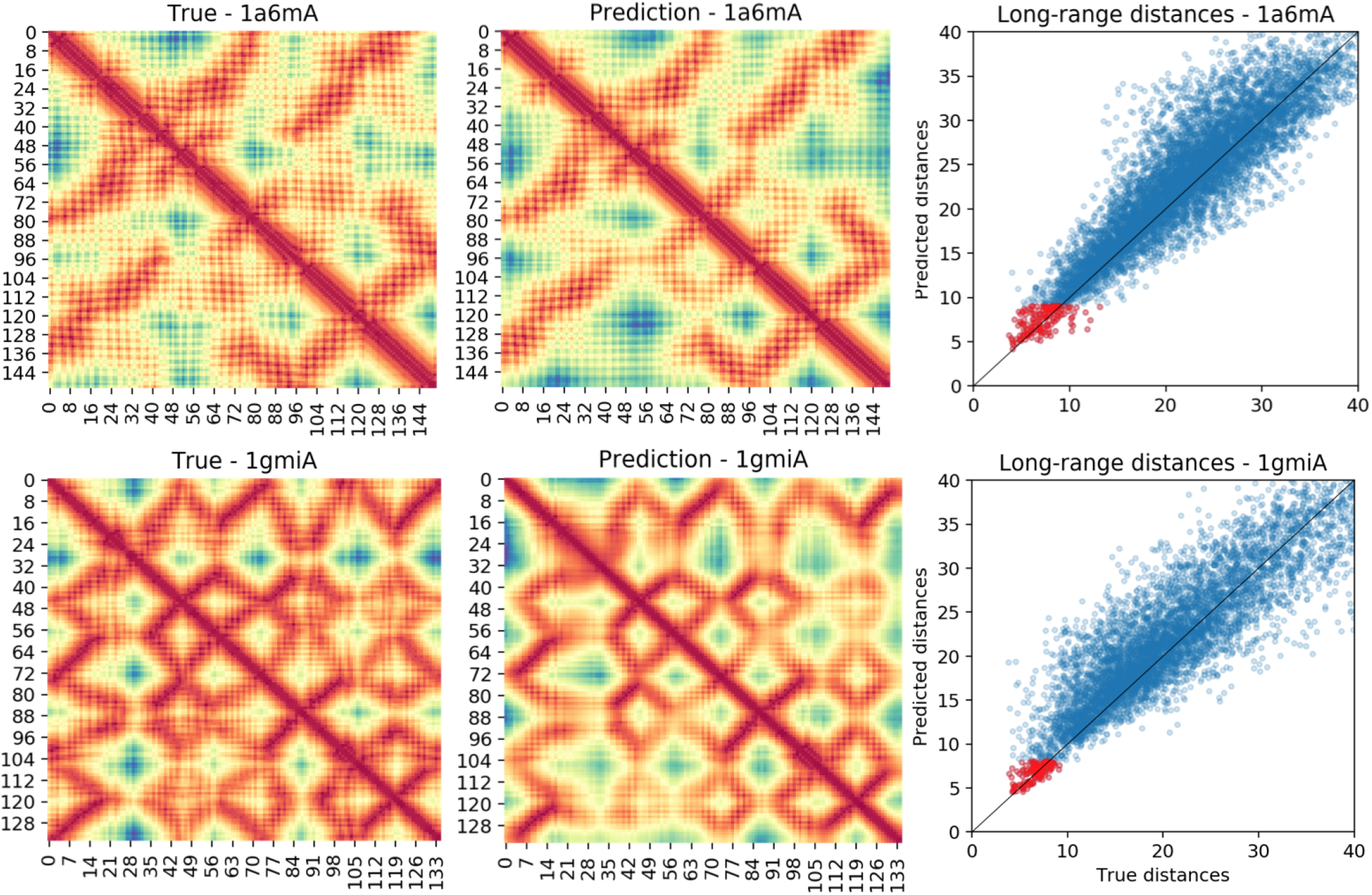
Comparison of true long-range distances and the distances predicted by our DL-Distance method for two random examples from the test dataset - ‘1a6m chain A’ (top row), ‘1gmi chain A’ (bottom row). True distance maps and predicted distance maps are shown in the first and second columns respectively. The plots in the last row visualize the comparison of predicted long-range distances and the corresponding true distances. Top L long-range predictions are shown in red and all other long-range distances are shown in blue.

### 3.4 Evaluation on CASP13 and CAMEO hard targets

We further evaluate our methods on two most difficult datasets - 131 hard targets in the CAMEO competition released after December 8, 2018, and the free-modeling targets of the CASP13 competition. Because of CASP’s agreements and policies, unlike the assessors of CASP13 and some of the top groups, we (public) do not have access to all the native structures. Of the 32 free-modeling domains, we only have access to 9 domains. Hence, we limit our evaluation to these 9 domains. Consistent with the practice, we predict distances and contacts for the full target and evaluate on the domains [24, 25]. For all our predictions, we use the alignments predicted by the trRosetta method, as our input. These alignment files are made publicly available by the Yang group at https://yanglab.nankai.edu.cn/. On these nine free-modeling domains our DL-Contact and DL-distance methods achieve *P_L_* of 38.8% and 32.3% compared to 45.0% by the RaptorX method, the top group in the competition (see Table 3). For evaluating our methods on the 131 CAMEO hard targets, we used the alignments generated using the same trRosetta method. Eight of these 131 targets’ native structure did not have any long-range contacts and so we skip them from our evaluation. When long-range contacts are evaluated, our results show that our DL-Contact and DL-Distance methods predict contacts with precision similar to the trRosetta method [16] (*P_L_* for trRosetta, DL-Contact, and DL-Distance are 48.0%, 48.3% and 46.7% respectively). Overall, this comparable performance of PDNET models with other state-of-the-art methods that use additional strategies such as using larger training sets and model ensembling, highlight the representativeness and value of PDNET.

**Table 3:** Summary of the performance of DL-Contact and DL-Distance on the 9 CASP13 free-modeling domains (for which the native structures are publicly available) and on the 131 CAMEO hard-targets. Performance of CASP13’s top group and the trRosetta method are provided for reference. Results of the trRosetta method are copied from the paper [16] since the predicted contacts are not publicly available.

| Dataset | Method | $P_L$ | $P_{NC}$ |
| --- | --- | --- | --- |
| CASP13 FM | DL-Contact | 38.8 | 38.5 |
|  | DL-Distance | 32.3 | 30.3 |
|  | Top CASP13 group | 45.0 | 45.7 |
| CAMEO-HARD | DL-Contact | 48.3 | 45.1 |
|  | DL-Distance | 46.7 | 43.7 |
|  | trRosetta | 48.0 | - |

### 3.5 Comparison with other datasets

Besides us, other groups have also attempted to ‘democratize’ the deep learning of protein distance prediction. The ProteinNet dataset by M. Alquraishi [29] is released as a standardized data set for machine learning of protein structure. It consists of large and small subsets for learning many features. Although the full dataset is 4 Terabytes in size, smaller subsets are available for download upon request. Benchmark results for contact prediction or distance prediction are not available for this dataset. Similarly, the Song group has released TAPE [30], a dataset packaged for predicting secondary structures, inter-residue contacts, and remote homology detection. These datasets can also be extended to predict inter-residue distances. Our work here, however, serves as the first work, not only to discuss the distance prediction problem as the primary focus but also to release a full deep learning framework to train and evaluate distance predictions.

## 4 CONCLUSION

We believe that PDNET will allow a faster progress for developing deep learning methods for protein structure prediction. It will be particularly helpful for researchers with some machine learning background interested in difficult problems like protein structure prediction. PDNET can be easily extended to test the significance of adding other features such as covariance matrix [14], precision matrix [8], and to predict dihedral angles/orientations [16]. We believe that significant future contributions can be made by focusing on novel feature engineering techniques, loss functions, and architectures that may be particularly suitable for this specific problem of distance prediction.

## 5 ACKNOWLEDGEMENTS

We thank Bikash Shrestha, Jacob Barger, Mrinal Rawool, David Richards, Amarilda Dyrmishi, Patrick Kong, and Anthony Ackah-Nyanzu at the University of Missouri-St. Louis for interesting discussions during this work. We are also extremely thankful to the reviewers of the International Conference on Machine Learning (ICML) 2020 conference for providing many useful comments.

## AVAILABILITY

All scripts, training data, deep learning code for training, validation, and testing, and Python notebooks are available at https://github.com/ba-lab/pdnet.

